# Glutamatergic facilitation of neural responses in MT enhances motion perception in humans

**DOI:** 10.1101/283994

**Authors:** Michael-Paul Schallmo, Rachel Millin, Alex M. Kale, Tamar Kolodny, Richard A.E. Edden, Raphael A. Bernier, Scott O. Murray

**Author notes:** corresponding author, F212/2C West Building, 2450 Riverside Ave S, Minneapolis, MN 55454. The authors declare no competing financial interests.

## Abstract

There is large individual variability in human neural responses and perceptual abilities. The factors that give rise to these individual differences, however, remain largely unknown. To examine these factors, we separately measured fMRI responses to moving gratings in the motion-selective region MT, and perceptual duration thresholds for motion direction discrimination within the same group of male and female subjects. Further, we acquired MR spectroscopy data that allowed us to quantify an index of neurotransmitter levels in the region surrounding MT. We show that individual differences in the Glx (glutamate + glutamine) signal in the MT region are associated with both higher fMRI responses and improved psychophysical task performance. Our results suggest that individual differences in baseline levels of glutamate within MT contribute to motion perception by increasing neural responses in this region.

**Significance:** What factors govern the relationship between neural activity and behavior? Our results suggest that one such factor is the level of glutamate, an excitatory neurotransmitter, within a particular region of cortex. By measuring an index of glutamate *in vivo* using magnetic resonance spectroscopy, we show that human subjects with more glutamate in the visual motion area known as MT also have larger fMRI responses (an index of neural activity) in this region. Further, people with more glutamate in MT can accurately perceive moving images presented more briefly within a behavioral task. Our findings point to an important role for glutamate levels in determining the relationship between neural responses and behavior during visual motion perception.

## Introduction

A direct relationship between greater neural responses and better perceptual functioning is well established in both humans (Boynton et al., 1999) and animal models (Newsome et al., 1989; Britten et al., 1992). One factor that may determine neural responsiveness and subsequent behavior is the amount of the neurotransmitter glutamate (Glu) available within a given region of cortex. Glu is the primary excitatory neurotransmitter in cortex and is released from presynaptic vesicles as the result of an action potential (Magistretti et al., 1999). Individuals with higher baseline Glu levels in a particular region may therefore possess greater potential for excitation within the local neural population (Conti and Weinberg, 1999). An index of Glu concentration can be measured non-invasively *in vivo* using MR spectroscopy (MRS). Although MEGA-PRESS sequences (Mescher et al., 1998) are most often used to measure γ-aminobutyric acid (GABA) levels, the difference spectrum that is obtained also contains a peak at 3.75 ppm associated with Glu. The size of this peak is believed to reflect the level of Glu within the MRS voxel, which is considered a stable individual trait. However, both Glutamine and Glutathione also contribute to the size of this peak (Mullins et al., 2014; Harris et al., 2017) – hence the peak is often referred to as Glx, to signify that it is a combined measure of multiple metabolites (glutamate, glutamine, & glutathione). Currently, it is not clear how Glx levels measured with MRS are related to the neural responses that support behavior.

We hypothesized that greater regional concentrations of Glx would be associated with higher neural activity (reflected in stronger fMRI responses) and in turn, superior performance on a task that depends on neural response magnitude. Our group recently examined the role of GABA during motion perception in humans using MRS (Schallmo et al., 2018). Here, we again chose to focus on neural processing within cortical area MT, in order to test our above hypothesis regarding a link between Glx, neural responses, and task performance. Neural responses in MT in both monkeys (Britten et al., 1992; Huk and Shadlen, 2005; Churan et al., 2008; Liu et al., 2016) and humans (Tootell et al., 1995; Rees et al., 2000; Tadin et al., 2011; Turkozer et al., 2016; Chen et al., 2017; Schallmo et al., 2018) are known to be tightly linked to motion perception. In particular, studies in humans suggest that motion duration thresholds (Tadin et al., 2003; Tadin, 2015) – the amount of time that a stimulus needs to be presented to accurately discriminate motion direction – are shorter under conditions that elicit higher MT responses (Tadin et al., 2011; Turkozer et al., 2016; Schallmo et al., 2018). Consistent with our hypothesis, we observed a link between individual differences in Glx levels and fMRI response magnitudes within human MT complex (hMT+): individuals with higher Glx had higher fMRI responses. Further, we found that both higher Glx and larger fMRI responses were associated with reduced motion duration thresholds (superior performance). Overall, our findings suggest that individual differences in the amount of Glu, as measured by MRS, contribute to motion direction discrimination by facilitating neural responses within hMT+.

## Methods

### Participants

Twenty-two young adults participated (mean age = 24 years, *SD* = 3.7; 13 females and 9 males). These subjects were included in two recent studies from our group examining the role of GABA in motion perception (Schallmo et al., 2018), and sex differences in motion processing (Murray et al., submitted). Subjects were screened for having normal or corrected-to-normal vision, no neurological impairments, and no recent psychotropic medication use. Further screening prior to MRS scanning included: no more than 1 cigarette per day in the past 3 months, no recreational drug use in the past month, no alcohol use within 3 days prior to scanning. Subjects provided written informed consent prior to participation and were compensated $20 per hour. All procedures were approved by the Institutional Review Board at the University of Washington (approval numbers 48946 & 00000556) and conformed to the guidelines for research on human subjects from the Declaration of Helsinki.

### Visual display and stimuli

For fMRI, stimuli were presented using either an Epson Powerlite 7250 or an Eiki LCXL100A (following a hardware failure), both with 60 Hz refresh rate. Images were presented on a screen at the back of the scanner bore and viewed through a mirror mounted on the head coil. Images were shown using Presentation software (Neurobehavioral Systems, Berkeley, CA). For psychophysics, a ViewSonic PF790 CRT monitor (120 Hz) was used with an associated Bits# stimulus processor (Cambridge Research Systems, Kent, UK). Stimuli were presented on Windows PCs in MATLAB (MathWorks, Natick, MA) using Psychtoolbox-3 (Brainard, 1997; Pelli, 1997; Kleiner et al., 2007). Viewing distance for both displays was 66 cm, and display luminance was linearized.

The visual stimuli were identical to those described previously (Schallmo et al., 2018). Briefly, drifting sinusoidal luminance modulation gratings were presented with Gaussian blurred edges on a mean gray background. Grating contrast was either 3% or 98%. Gratings were 2° in diameter for fMRI, and 0.84° in diameter for psychophysics. Spatial frequency was 1 cycle/° (fMRI) or 1.2 cycles/° (psychophysics). Drift rate was 4 cycles/s for both experiments.

### Experimental procedure and data analysis

#### Functional MRI

The fMRI paradigm has been described previously (Schallmo et al., 2018). Structural (1 mm resolution) and functional data (3 mm resolution, 30 oblique-axial slices, 0.5 mm gap, 2 s TR) were acquired on a Philips 3T scanner. Before the fMRI task scans, a 1-TR scan was acquired with the opposite phase-encode direction, which was used for distortion compensation.

The fMRI task measured the response to drifting gratings presented at different contrast levels within a blocked experimental design (Figure 1A). Sixteen gratings were presented within each block (400 ms duration, 225 ms inter-stimulus interval, 10 s total block duration), which drifted in 8 possible directions (randomized & counterbalanced). Stimulus contrast varied across block; this began with a 0% contrast (blank) block. Then, blocks of high (98%) and low contrast (3%) gratings were presented in alternating order, each followed by a blank block to allow the fMRI response to return to baseline (6 high, 6 low, and 13 blank blocks total). Task scans were 4.2 minutes long (125 TRs), and each subject completed a total of 2-4 scans across 1 or 2 scanning sessions, as part of a larger set of visual fMRI experiments.

**Figure 1.**
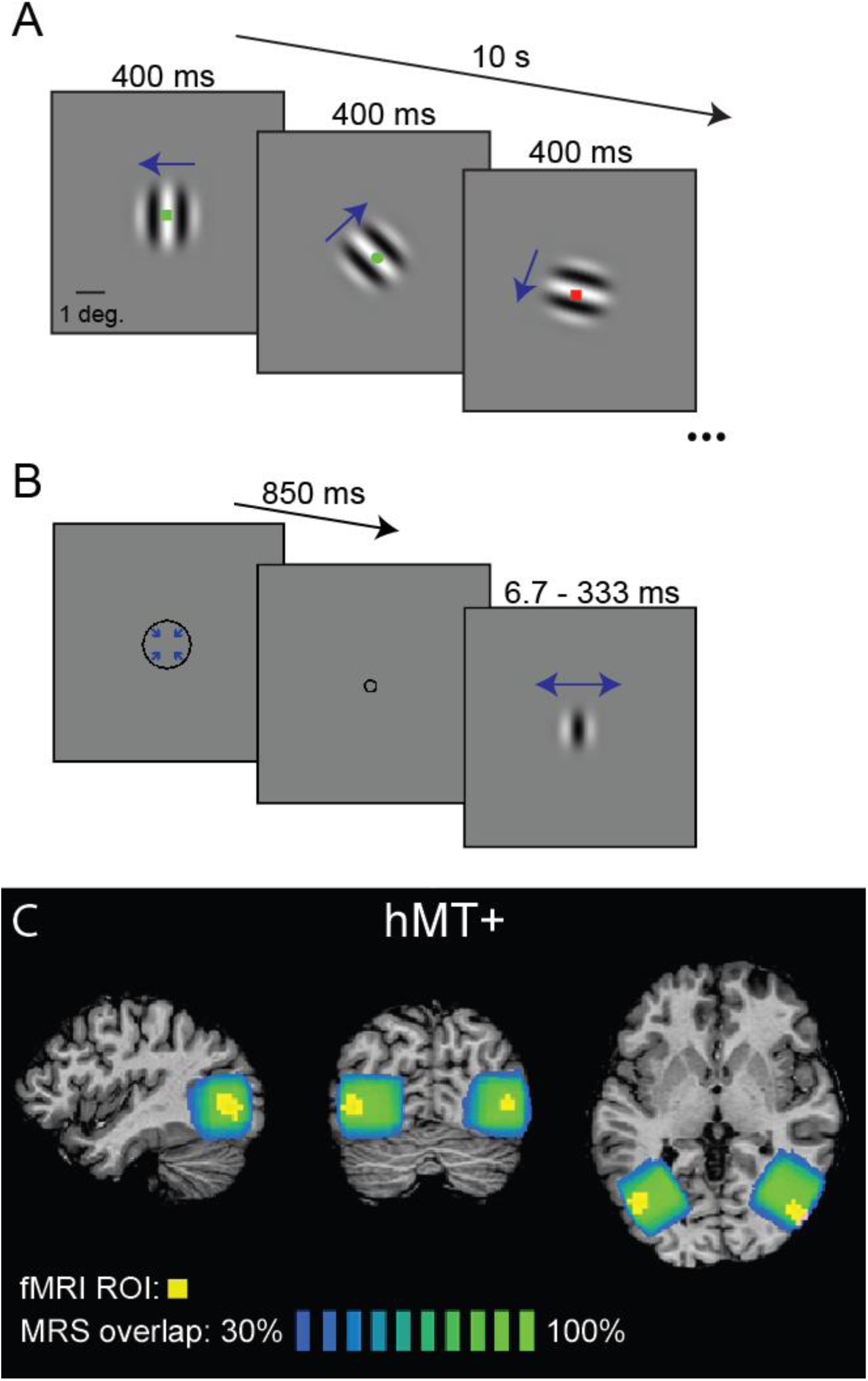
Visual stimuli and MR spectroscopy. **A** shows the fMRI task timing (10 s blocks of 400 ms drifting gratings). Blue arrows indicate motion direction. Fixation task stimuli also shown. **B** shows the timing of a psychophysics trial from the task performed outside the scanner (850 ms cue, variable grating duration). Average MRS voxel placement is shown in **C** (adapted from Schallmo et al., 2018). Green-blue color indicates the percent overlap of the fMRI-localized MRS voxels in the hMT+ region (in Talairach space) across all subjects. Average hMT+ ROIs from fMRI for all subjects are shown in yellow (threshold correlation between predicted & observed fMRI timeseries *r* ≥ 0.3).

A functional localizer scan was also included in each scanning session, in order to identify regions-of-interest (ROIs). This localizer was designed to identify human MT complex (hMT+); we did not attempt to distinguish areas MT and MST (Huk et al., 2002). Drifting gratings (as above, but 15% contrast) alternated with static gratings across blocks (10 s block duration, 125 TRs total). Bilateral hMT+ ROIs (averaged across subjects, in Talairach space) are shown in Figure 1C (yellow).

Subjects performed a fixation task during all fMRI scans. This involved responding to a green circle within a series of colored shapes presented briefly at fixation (Figure 1A). This task encouraged subjects to focus their spatial attention at the center of the screen and minimized eye movements away from this position.

FMRI data were analyzed using BrainVoyager (Brain Innovation, Maastricht, The Netherlands), and MATLAB (The Mathworks, Natick, MA). Preprocessing was performed in the following order: motion correction, distortion compensation, high-pass filtering (2 cycles/scan), co-registration, and transformation into the space of the individual subject’s T_1_ anatomical scan. Normalization and spatial smoothing were not performed; all analyses were conducted within ROIs for each individual subject. ROIs were defined using correlational analyses, as previously described (Schallmo et al., 2018). ROI position was verified on an inflated model of the white matter surface. Analyses of fMRI time course data was performed in MATLAB using BVQXTools. Data were broken into epochs spanning 4 s before the onset of each block to 2 s after the offset. Average baseline responses were calculated across blocks separately for each condition (3% & 98% contrast); this was computed as the average response to the blank background 0-4 s prior to block onset. Data were converted to percent signal change, averaged across blocks, across ROIs from each hemisphere, and then averaged across scanning sessions. Responses for each condition were defined as the mean of the fMRI signal peak (from 8-12 s following block onset). For the correlational analyses (i.e., Figure 2C), fMRI responses to low and high contrast stimuli were averaged, as an index of overall responsiveness.

**Figure 2.**
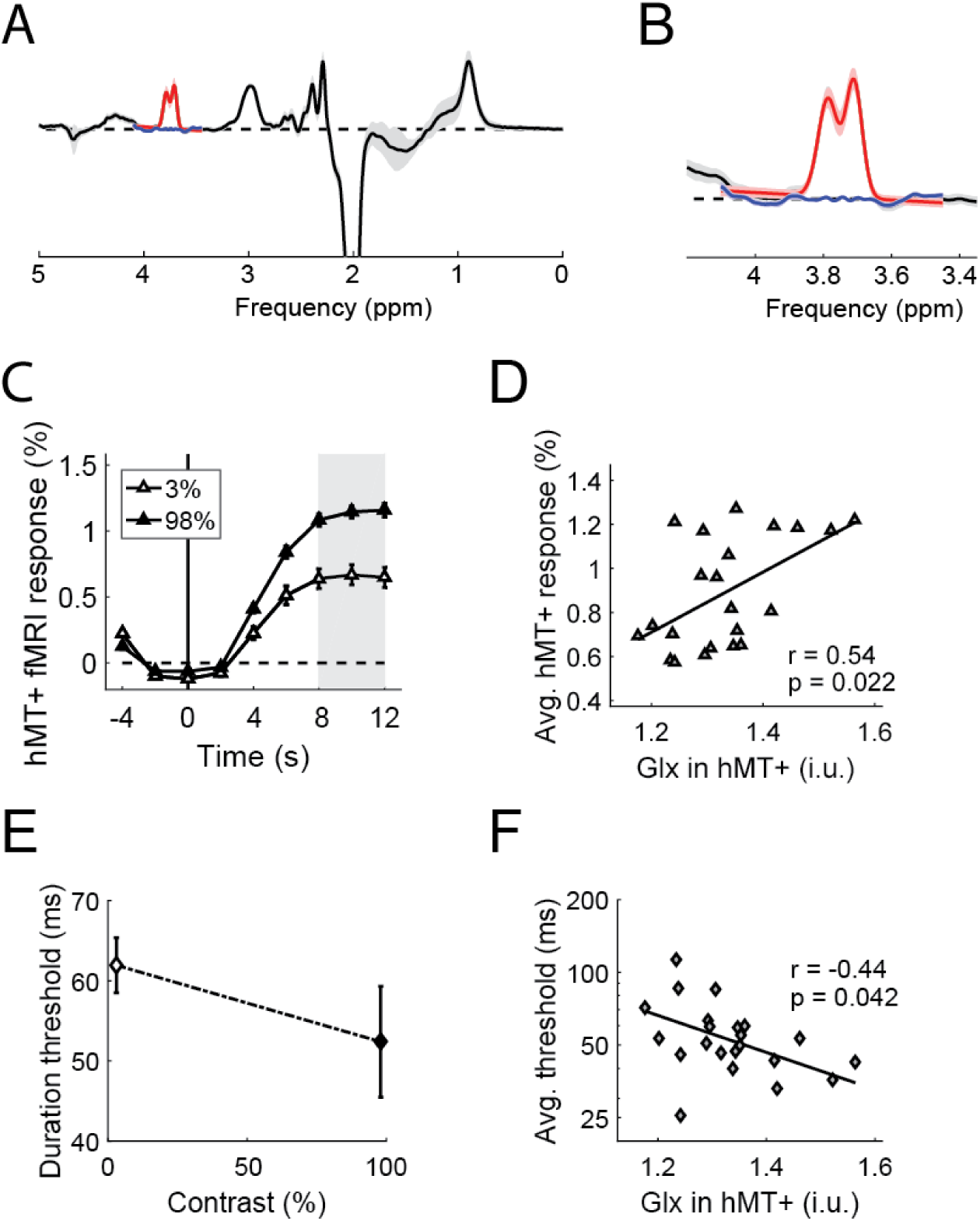
A link between Glx, fMRI, and motion duration thresholds. **A** shows the MR spectra (black), fits to the Glx peak (red) and residuals (blue) averaged across subjects. Error bars in **A** & **B** show *SD*. **B** shows a zoomed-in view of the data from **A.** Panel **C** shows the time course of fMRI responses measured in hMT+, averaged across subjects. Peak response was calculated within the gray region. Higher Glx in hMT+ correlates significantly with greater fMRI responses in the same area (**D**; averaged across contrasts). Duration thresholds are shown in **E**; these correlated negatively with Glx in hMT+ (**F**; geometric mean of thresholds across contrasts). Error bars in **C** & **E** show *SEM*.

#### Behavioral psychophysics

Subjects performed a psychophysical motion direction discrimination task outside of the scanner, during a separate experimental session. This paradigm followed established methods (Tadin et al., 2003; Foss-Feig et al., 2013) and is described in a recent paper from our group (Schallmo et al., 2018). In the full experiment, drifting gratings were presented at 2 contrast levels (3% & 98%) and 3 sizes (0.84, 1.7, & 10° diameter). In order to minimize the effects of spatial suppression / summation seen with larger stimuli (Tadin et al., 2003; Foss-Feig et al., 2013; Tadin, 2015; Schallmo et al., 2018), only data from the smallest stimulus size were included in the current study. Thus, we focused on the smallest size condition, wherein motion discrimination performance is expected to depend primarily on the magnitude of neural responses driven by stimuli within the classical receptive field (Tadin, 2015).

In this task, subjects were asked to report whether a briefly presented grating drifted left or right (Figure 1B). The stimulus duration was adjusted across trials (range 6.7 – 333 ms) using the Psi adaptive staircase method within the Palamedes toolbox (Prins and Kingdom, 2009; Kingdom and Prins, 2010), in order to find the duration for which motion discrimination performance reached 80% accuracy. Trials began with a brief central fixation mark (a shrinking circle; 850 ms), followed by the presentation of the grating. A fixation mark presented after the grating cued the subjects to respond; response time was not limited. Staircases were comprised of 30 trials for each stimulus condition (3% & 98% contrast), presented in a randomized counterbalanced order within each run. Run duration was approximately 6 min. Subjects completed 4 runs each, for a total experiment duration of about 30 min, including instructions and practice prior to the start of the main experiment.

Data from each staircase were fit with separate Weibull functions using the Palamedes toolbox (Prins and Kingdom, 2009). Motion duration thresholds were defined at 80% accuracy based on this fit. When averaging across thresholds from different conditions (i.e., for the correlations in Figure 2D), the mean threshold was first taken across runs in each condition, and then the geometric mean was taken across conditions, to account for the fact that threshold ranges varied across conditions.

#### MR spectroscopy

Our MRS data acquisition was the same as described previously (Schallmo et al., 2018). Briefly, we used a MEGA-PRESS (Mescher et al., 1998) sequence on a Philips 3T scanner. Sequence parameters were as follows: 3 cm isotropic voxel, 320 averages, 2048 data points, 2 kHz spectral width, 1.4 kHz bandwidth refocusing pulse, VAPOR water suppression, 2 s TR, 68 ms TE, 14 ms editing pulses at 1.9 ppm for “on” and 7.5 ppm for “off” acquisitions, “on” and “off” interleaved every 2 TRs with a 16-step phase cycle. Across 2 additional scanning sessions (separate from fMRI), MRS data were acquired from the following 3 regions: 1) lateral occipital cortex, centered on hMT+ (Figure 1C); 2) mid-occipital region of early visual cortex (EVC), parallel to the cerebellar tentorium; 3) fronto-parietal cortex (FPC), centered on the “hand-knob” (Yousry et al., 1997).

Voxels in hMT+ were positioned using an on-line functional localizer (moving vs. static, as above); left and right hMT+ was measured for each subject, typically within the same scanning session. For EVC and FPC, voxels were positioned based on each individual’s anatomical landmarks; see (Schallmo et al., 2018) for images of these voxel positions. Data were acquired in EVC across 2 separate scans (typically in different sessions; 1 subject had only 1 EVC scan). Only 1 scan was acquired in FPC (1 subject did not complete the FPC scan). Subjects watched a film of their choice during MRS, with images presented on the screen (as above, for fMRI), and audio presented through MR-compatible headphones. MRS, fMRI, and psychophysical data were collected within a 2-week period for all subjects. Previous work suggests metabolite measurements from MRS are stable over the course of several days (Evans et al., 2010; Greenhouse et al., 2016).

MRS data were analyzed using the Gannet toolbox version 2.0 (Edden et al., 2014). Automated processing steps within this toolbox were as follows: frequency and phase correction, artifact rejection (frequency correction > 3 SD above the mean), and exponential line broadening (3 Hz). The Glx peak at 3.75 ppm was fit with a double-Gaussian function (Figure 2A & B); the integral of this function served as the measure of Glx concentration. Note that this fitting was performed separately from fitting the GABA peak at 3 ppm. The value of Glx was scaled by the integral of the water peak for each individual. Correction for gray / white matter content within the voxel was not performed.

No significant correlations were observed between the concentration of unsuppressed water (fit by Gannet) in hMT+ and either fMRI response magnitudes (*r*_20_ = −0.37, FDR corrected 2-tailed *p* = 0.081) or psychophysical thresholds (*r*_20_ = 0.44, FDR corrected 2-tailed *p* = 0.080). However, as these correlations were moderately strong (despite not reaching statistical significance), the effect of scaling Glx values to water merits further consideration. Since we observed qualitatively similar results to those presented below when scaling Glx to creatine instead of water (not shown), we believe it is reasonable to conclude that the observed correlations are driven by a genuine relationship between Glx, fMRI, and motion thresholds, rather than being attributable to water scaling.

### Statistics

Statistical analyses were performed in MATLAB. Because variance differed for duration thresholds at low and high contrast, Friedman’s non-parametric ANOVA was used for this comparison. One tailed Pearson’s correlation coefficients were calculated for all correlational analyses; this is justified by the strong *a priori* hypotheses being tested and is noted for each occurrence in the Results. Correlation significance was further examined using (non-parametric) permutation tests, which involved randomly shuffling the data being correlated across subjects in each of 10,000 iterations. The proportion of permuted correlations with coefficients greater (or more negative) than that of the observed, un-shuffled correlation served as the measure of significance (the *p*-value).

## Results

Using MR spectroscopy, we measured the concentration of glutamate (plus co-edited metabolites; Glx; Figure 2A & B) within the motion selective region of human MT complex (hMT+). We sought to determine whether individual differences in Glx concentration were positively associated with neural responsiveness during motion perception, as indexed with fMRI. FMRI signals were measured in hMT+ in response to moving gratings with low (3%) and high (98%) luminance contrast (Figure 2C). Responses were larger for high than for low contrast stimuli (1-tailed paired *t*_21_ = 8.02, *p* = 4 × 10^-8^), as expected (Tootell et al., 1995). An overall index of fMRI responsiveness was obtained for each subject by averaging the peak response to both low and high contrast (gray regions; Figure 2C). As predicted, there was a significant positive correlation between average fMRI responses in hMT+ and concentrations of Glx in the same region (Figure 2D; *r*_20_ = 0.54, 1-tailed *p* = 0.022; FDR corrected). These results suggest that individuals with more Glx in the region surrounding area MT have larger neural responses to moving stimuli, consistent with stronger glutamatergic excitation.

We next examined whether individual differences in Glx concentration in the region surrounding hMT+ were associated with behavioral differences in motion discrimination. We measured motion duration thresholds (Tadin et al., 2003; Tadin, 2015) to small (0.84°) gratings of low and high contrast (Figure 2E). Duration thresholds were smaller for high vs. low contrast stimuli (Friedman’s *χ*^2^_1_ = 30, *p* = 4 × 10^-8^), consistent with previous findings (Tadin et al., 2003; Foss-Feig et al., 2013). To obtain an overall measure of motion perception, duration thresholds were averaged across low and high contrast. We observed the predicted relationship between Glx in hMT+ and average motion duration thresholds; individuals with greater Glx showed lower thresholds (superior performance; Figure 2F; *r*_20_ = −0.44, 1-tailed *p* = 0.042; FDR corrected). The relationship between duration thresholds and Glx was specific to hMT+; we saw no significant correlations between thresholds and Glx in two other MRS voxels (early visual cortex [EVC] and fronto-parietal cortex; |*r*_20_| < 0.23, uncorrected *p*-values > 0.14).

The significant positive correlation between Glx and fMRI response magnitudes, and the negative correlation between Glx and duration thresholds together suggest a negative relationship may exist between fMRI responses and duration thresholds. Indeed, we observed a significant negative correlation (*r*_20_ = −0.60, 1-tailed *p* = 5 × 10^-4^); higher averaged fMRI responses in hMT+ were associated with lower averaged duration thresholds, consistent with our observations from a separate analysis of different stimulus conditions within the same dataset (Murray et al., submitted). Taken together, our findings are consistent with the idea that stronger glutamatergic excitation drives larger neural responses in hMT+ during motion perception, thereby facilitating lower duration thresholds (i.e., superior motion discrimination).

The concentration of Glx showed some specificity between regions. Glx concentrations in hMT+ showed some association with Glx in EVC, but this did not survive correction for multiple comparisons (*r*_20_ = 0.43, 1-tailed *p* = 0.023, uncorrected; *p* = 0.069 after FDR correction). Neither hMT+ nor EVC concentrations were significantly associated with Glx in fronto-parietal cortex (|*r*_20_| < 0.37, FDR corrected 1-tailed *p*-values > 0.097).

Recent work from our group has demonstrated a small but statistically reliable gender difference in motion discrimination thresholds, with females showing slightly higher duration thresholds on average than males (Murray et al., submitted). Thus, we examined whether there might be gender differences in Glx levels in hMT+ as well. However, no significant difference in hMT+ Glx was observed between males (mean = 1.33 i.u., *SD* = 0.10) and females (mean = 1.34 i.u., *SD* = 0.10; 2-tailed independent samples *t*_20_ = 0.17, *p* = 0.87). This is concordant with our recent finding (Murray et al., submitted) that fMRI response magnitudes in hMT+ also did not differ between genders (despite the significant difference in psychophysical thresholds for the same subjects). The lack of a gender difference in Glx may reflect the fact that, although neural processing in MT clearly influences motion perception, response magnitudes in MT are not the only factor that determine duration thresholds for motion discrimination.

We also examined whether the observed relationships between Glx and fMRI or motion discrimination thresholds might in fact be attributable to GABA levels, possibly as an artifact of the MRS sequence. Although a Glx peak is obtained using MEGA-PRESS, this sequence is typically used to measure the concentration of an edited GABA peak at 3 ppm, which is acquired in the same scan (Mescher et al., 1998; Mullins et al., 2014). We have previously observed that higher GABA concentrations in hMT+ correlate with lower motion discrimination thresholds within the same group of subjects (Schallmo et al., 2018). Thus, we sought to determine whether the current relationships with Glx might be accounted for by the previously reported relationship with GABA – perhaps due to homeostatic processes balancing the levels of these neurotransmitters, the manner in which they were measured together using MEGA-PRESS, and/or the method of quantifying both peaks in Gannet (Edden et al., 2014). However, we observed no significant correlations between Glx and GABA measurements across individuals in any of the 3 ROIs that we examined (not shown; all |*r*_(20)_| < 0.34, 2-tailed *p*-values > 0.14; uncorrected), suggesting that the Glx results presented here cannot be explained by the amount of co-measured GABA.

## Discussion

To our knowledge, this is the first study to present evidence of a 3-way link between behavioral performance, BOLD response magnitudes, and Glx levels in humans. These findings help to clarify the role of glutamate in visual motion perception, and suggest that higher excitatory tone (as measured by Glx from MRS) facilitates larger neural responses and greater perceptual sensitivity. A straightforward association between higher levels of Glx, larger fMRI responses, and superior performance is perhaps not surprising. However, our findings are notable, given that the relationships between each of these measures and the underlying neural responses are complex (Logothetis, 2008; Duncan et al., 2014; Harris et al., 2015). Adding to this complexity, Glu also plays an important role in cell metabolism (Magistretti et al., 1999) in addition to functioning as a neurotransmitter.

It is still unclear how measures of Glx from MRS are tied to changes in neural activity in the human brain (Duncan et al., 2014). Studies using functional MRS at 7 Tesla to measure Glu show increased occipital Glu following visual stimulation, and support the idea that the magnitude of Glu being measured depends on the level of local neural activity (Mangia et al., 2007; Lin et al., 2012; Schaller et al., 2013; Apšvalka et al., 2015). However, it is not yet known to what extent the Glu being measured with MRS reflects a ‘direct’ relationship with neural activity (i.e., driven by Glu neurotransmission) vs. an ‘indirect’ one (i.e., reflects Glu’s role in cell metabolism, which is affected by spike rate). It has been argued that Glu becomes more MR visible as it moves from pre-synaptic vesicles to the synaptic cleft during neurotransmission and into astrocytes following reuptake, and that this change in MR visibility may suggest that Glx measured with MRS reflects Glu release during neurotransmission (Apšvalka et al., 2015). Alternatively, higher rates of neural activity might also increase the rate of Glu cycling through the synaptic cleft, which could lead to a transient buildup of Glu (depending on the rate limiting step in this cycle; Lin et al., 2012). While the 3-way association between Glx, fMRI, and behavior in the current study is consistent with the idea that Glx levels reflect the strength of glutamatergic neurotransmission, direct experimental support for this hypothesis remains lacking.

Our work helps to clarify how glutamate in human visual cortex supports visual behavior. We are aware of very few studies examining how visual perception is related to individual differences in Glx measurements. Some have found that higher occipital Glx measurements are associated with increased visual functioning (Terhune et al., 2015; Wijtenburg et al., 2017), while others have not (Pugh et al., 2014; Takeuchi et al., 2017). One study from the latter category found that frontal Glx levels, but not measurements in MT, were associated with individual differences in an ambiguous motion perception task (Takeuchi et al., 2017). The discrepancy with the current findings may be explained by the manner in which visual behavior was assessed (i.e., ambiguous motion vs. direction discrimination). The association between lower motion duration thresholds (better performance) and greater Glx in MT from the current study suggests that motion discrimination performance is facilitated by higher levels of Glu, likely due to greater excitatory neural activity within area MT.

The role of area MT in visual motion perception is well established (Born and Bradley, 2005; Zeki, 2015), but the manner in which individual differences in MT responsiveness relate to differences in perception is less clear. In a seminal study, Rees and colleagues (Rees et al., 2000) demonstrated a positive linear relationship between the coherence of moving dot stimuli and the fMRI response in human MT. They also reported modest individual variability in the coherence-response function, but not whether such variability corresponds to differences in perception. A more recent study has found a correspondence between improved behavioral performance and fMRI response changes in MT following perceptual learning (Chen et al., 2017). Specifically, subjects with smaller thresholds in a motion discrimination paradigm following extensive training also showed greater sharpening of tuning within MT, as assessed by multi-voxel pattern analysis. This suggests that increased selectivity within MT is important for learning to perform a motion discrimination task.

Here, we used a variation of a well-established motion discrimination paradigm (Tadin et al., 2003; Foss-Feig et al., 2013; Tadin, 2015), in which larger neural responses (particularly within area MT) are thought to facilitate motion direction discrimination for more-briefly presented stimuli, resulting in shorter duration thresholds (i.e., better performance). Evidence for the role of neural activity in MT within this paradigm has been provided by studies in both macaques (Liu et al., 2016) and humans (Tadin et al., 2011; Turkozer et al., 2016; Schallmo et al., 2018). Our current findings build upon this work and suggest that individual differences in Glu concentration in area MT contribute to the neural responsiveness within the region, as well as consequent motion discrimination performance. Measuring Glu may thus provide valuable insight into the neural basis of individual differences in visual perception.

## Acknowledgments

We thank Anastasia V. Flevaris for help with data acquisition and analysis. We also thank Brenna Boyd, Judy Han, Heena Panjwani, Micah Pepper, Meaghan Thompson, Anne Wolken, and the UW Diagnostic Imaging Center for help with subject recruitment and/or data collection. This work was supported by funding from the National Institute of Health (F32 EY025121 to MPS, R01 MH106520 to SOM, T32 EY00703). This work applies tools developed under NIH grants R01 EB016089 and P41 EB015909; RAEE also receives support from these grants.

Current affiliation for MPS: Department of Psychiatry, University of Minnesota, Minneapolis, MN

